# *In vitro* analysis of colistin and ciprofloxacin antagonism of *Pseudomonas aeruginosa* phage PEV2 infection activities

**DOI:** 10.1101/2020.12.02.406561

**Authors:** Katarzyna Danis-Wlodarczyk, Alice Cai, Anna Chen, Marissa Gittrich, Matthew B. Sullivan, Daniel J. Wozniak, Stephen T. Abedon

## Abstract

Phage therapy is a century-old technique employing viruses (phages) to treat bacterial infections. In the clinic, phage therapy often is used in combination with antibiotics. Antibiotics, however, interfere with critical bacterial activities, such as DNA and protein synthesis, which also are required for phage infection processes. Resulting antagonistic impacts of antibiotics on phages nevertheless are not commonly determined in association with phage therapy studies using standard, planktonic approaches. Here we assess the antagonistic impact of two antibiotics, colistin and ciprofloxacin, on the bactericidal, bacteriolytic, and new virion production activities of *Pseudomonas aeruginosa* podovirus PEV2, using a broth culture, optical density-based ‘lysis profile’ assay. Though phage-antibiotic combinations were more potent in reducing cell viability than phages or antibiotics alone, colistin substantially interfered with phage PEV2 bacteriolytic and virion-production activities at minimum inhibitory concentration (MIC). Ciprofloxacin, by contrast, had no such impact at 1x MIC or 3x MIC. At higher but still clinically relevant concentrations (9× MIC) burst sizes were still significant (~30 phages/infected bacterium). We corroborated these lysis profile results by more traditional measurements (colony forming units, plaque forming units, one-step growth experiments) and two other *P. aeruginosa* phages. To our knowledge this is the first study in which detailed antibiotic impact on *P. aeruginosa* phage infection activities has been determined under conditions similar to those used to determine antibiotic MICs and could point especially to ciprofloxacin as a minimally antagonistic phage therapy co-treatment of *P. aeruginosa* infections.

## Introduction

“It’s not about phages *or* antibiotics. It’s about phages *and* antibiotics.” –Paul Grint^1^ Development of the first antibacterial chemotherapeutics, Salvarsan^2,3^, Prontosil^4^, and penicillin^5^, transformed clinical treatments of bacterial infections. Currently, antibiotic therapy represents the standard of care for treatment of these infections though its utility is threatened by an emergence of multi/pan-drug resistant or tolerant pathogens^6,7^. This has led to increased health care expenses and economic damages that are comparable to those of the 2008-2009 global financial crisis^8^.

The U.S. Centers for Disease Control and Prevention (CDC) estimates that more than 2.8 million antibiotic-resistant infections occur each year, leading to more than 35,000 deaths just in the United States^9^. Globally, this number increases to at least 700,000 deaths each year^8^ and is predicted to lead to 10 million deaths per year by 2050^10^. This is particularly concerning given the limited number of new antimicrobial agents, both currently available and in drug development pipelines, that are active especially against Gram-negative pathogens^11^. Moreover, infections by even antibiotic-sensitive bacteria can be tolerant of antibiotic treatments^12–18^. Thus, additional antibacterial agents and approaches are needed, with traits of interest including an ability to treat antibiotic-recalcitrant bacterial infections and reduce the potential for bacteria to evolve resistance.

One emerging antibacterial approach is bacteriophage therapy^19,20^. Bacteriophages (phages) can lyse and kill bacteria and collectively are the most abundant replicative entities in the biosphere^21^. They have been in common clinical use for decades in a number of countries, particularly those from former Soviet Union (Georgia and Russia) and Poland. Currently, in Europe as well as in the U.S.^22–27^, new phage therapy centers are opening and several new clinical trials taking place^28–33^.

Phage therapy in the clinic is often tested and used in combination with antibiotic treatments^24,34–45^. Crucial though with combination therapies is avoidance of substantial antagonisms between agents^46,47^. For phage therapy as typically practiced, this means that phages while infecting antibiotic-treated bacterial populations ideally should retain bactericidal, bacteriolytic, and virion production activities. It can be useful, therefore, to identify possible incompatibilities between specific phages and specific antibiotics early during the development of combination therapies, or in the course of compassionate use^20,25,27^. There has been comparatively little analysis, however, of the antagonistic impacts of antibiotics on the ability of phages to successfully infect antibiotic-sensitive bacteria, especially under standard, pharmacologically relevant conditions.

To assess the antagonistic impact of antibiotics on phage antibacterial activities, given the substantial numbers of possible combinations of therapeutic phage and antibiotic types, higher-throughput approaches are needed—ideally performed under conditions that are similar to those employed to determine antibiotic minimum inhibitory concentrations^48–50^. Here we employ a 96-well microtiter plate-compatible turbidimetric, ‘lysis profile’^51^ means of assessing phage infection activity in the presence and absence of antibiotics, as corroborated by more traditional broth-based methods. We find that colistin even at low bacteria-inhibiting concentrations is nearly fully antagonistic to *Pseudomonas aeruginosa* phage PEV2^52,53^ infection activities in broth. Antagonism by ciprofloxacin, in contrast, is much lower, with phages retaining substantial phage infection activities even at high, clinically relevant concentrations.

## Materials and Methods

### Bacteria, antibiotics, and phages

*P. aeruginosa* PAO1 Krylov is used as a reference strain in this study, as provided by Dr. Jean-Paul Pirnay (Queen Astrid Military Hospital, Brussels, Belgium). *P. aeruginosa* surface mutants used for phage receptor analysis are described in Table S1 and are isogenic to the *P. aeruginosa* PAO1 strain from Harvard University. Mueller-Hinton Broth II (MHB II), cation-adjusted (Becton Dickinson, USA), is used for bacteria liquid cultures, supplemented where indicated with different concentrations of ciprofloxacin (ciprofloxacin hydrochloride, usp reference standard, USA) or colistin (colistin sulfate, usp reference standard, USA). Minimal inhibitory concentrations (MIC) were measured by exposing a bacterial culture to increasing concentrations of antibiotic in MHB II, as previously described^54^. MIC for ciprofloxacin was determined as 0.91 μM = 303.2 ng/ml, and for colistin MIC was found to be 39.06 μM = 68.75 μg/ml (see Appendix A for further discussion). Antibiotic concentrations in basic experiments were varied as 0× (no antibiotic), 1×, 3×, 9×, 27×, and 81× MIC (× = times). Additional antibiotic concentrations were tested for ciprofloxacin between 9× and 27× MIC toward ascertaining maximal antibiotic concentrations that will still allow for phage bacteriolytic and virion production activities. We also confirmed that *P. aeruginosa* PA01 failed to replicate at 1× MIC colistin and ciprofloxacin using microscopic total count determinations (data not shown).

*P. aeruginosa* phage PEV2 (NC_031063.1, family *Podoviridae*, subfamily *Sepvirinae*, genus *Litunavirus*) was kindly provided by Elisabeth Kutter (Evergreen State College, USA). The phage was propagated in LB (Lysogeny Broth, Fisher Scientific, USA), purified via PEG precipitation (25% polyethylene glycol 8000, VWR, USA), subsequently suspended in phage buffer (10 mM Tris-HCl, Sigma,10 mM MgSO_4_, 150 mM NaCl, pH 7.5, Fisher Scientific), and stored at 4°C, as previously described^55^. Phage titers were assessed using the double-agar layer method^56,57^ and defined as plaque forming units per ml (PFU/ml).

Bacterial densities were estimated as colony forming units per ml (CFU/ml). When CFUs were determined under experimental conditions, e.g., as in the presence of phages, cultures were first exposed, prior to plating, to virucide consisting of 7.5% black tea infusion (Ceylon tea from Sri Lanka, Ahmad Tea, UK) and 0.53% FeSO_4_ (Sigma, Germany), prepared according to the Chibeu and Balamurugan^58^ protocol. The virucide was used to inactive phage virions to avoid overestimation of phage-mediated bacterial killing (bactericidal activity) prior to plating. In separate experiments this treatment was found not to affect bacterial viable counts (data not presented).

### Phage surface receptor analysis

Phage PEV2 specificity to particular bacterial receptor was tested on *P. aeruginosa* PAO1 mutants deficient in biosynthesis of flagella, type IV pili, alginate production or structure of their lipopolysaccharide (LPS) was modified (Table S1), as previously described^59^. Bacterial susceptibility to phage PEV2 was identified by spot testing (10 μl volume) using a suspension of 10^7^ PFU/ml. The plates were checked after 4–6h and again after 18h for the presence of a spot corresponding to confluent lysis located beneath applied volumes.

### Analysis of phage infection activity via lysis profiles

Lysis profiles involve the addition of phages to relatively high densities of bacteria, with phage impact on bacteria observed as reductions in bacterial culture turbidity. To assure robust measure of culture turbidity declines, i.e., as is generally assumed in these experiments to be associated with phage-induced bacterial lysis, cultures are not diluted following phage adsorption (contrast to one-step growth experiments, as described in the following section, where cultures instead are diluted following phage adsorption). The turbidity measurements were conducted at OD_550nm_ with the use of SpectraMax i3x Multi-mode Microplate Reader (Molecular Devices, USA).

Phage PEV2, antibiotic, or both were added to log-phase *P. aeruginosa* PAO1 (OD_600nm_ = 0.32, ~2.0 × 10^8^ CFU/ml; time, *t* = 0 min). Subsequent incubations were of at least 2h in length and took place as 200 μl/well volumes within 96-well microtiter plates. For incubations, these plates were covered with Breathe-Easy film (Diversified Biotech, USA) and kept at 37°C. Turbidity measurements were taken every 5 min and cultures were shaken for 3 sec before every read. All materials including microtiter plates were preincubated at experimental temperatures to limit physiological stress, and phages as well as antibiotics were prepared in MHB II prior to use. Negative controls consisted of untreated PAO1 and MHB II only. Input multiplicities of infection (MOIs)^60,61^ ranged from 0.1 (for virion production assessment) to 10 (for phage-induced bacteriolysis assessment). These MOIs corresponded to within-well titers of ~1 × 10^7^, ~1 × 10^8^, ~5.0 × 10^8^, and 1 × 10^9^ PFU/ml (or MOIs of 0.1, 1, 5, and 10, respectively). Following lysis profile determinations, PFU and CFU counts were assessed initially via spot test assays^56,57^ and for PFUs further via the double-layer method^56,57^. Experiments were performed in three technical repeats and three biological repeats.

### Analysis of phage infection activity via one-step growth

One-step growth experiments were undertaken according to previously established methods^62,63^, with modifications. A volume of 900 μl of a mid-exponential bacterial culture (final concentration ~2 × 10^8^ CFU/ml) in MHB II was mixed with 100 μl of PEV2 phage suspension (final concentration ~1.0 × 10^6^ PFU/ml) to obtain an input MOI of 0.005. Phages are allowed to adsorb for 8 min at 37°C, after which time the mixture is diluted 10^4^ fold, with samples then taken at 20-30 seconds intervals for titer determination. PFU/ml titers presented in the graphs are values from which unabsorbed phages values were subtracted. Antibiotic, if present, was added to PAO1 cultures 15 min prior to phage addition as well as to all dilution media. At least three biological repeats were performed.

### Statistics

Area under the curve (AUC) was calculated by GraphPad Prism version 7.00 for Windows (GraphPad Software, La Jolla California USA, www.graphpad.com) based on 5h lysis profiles. Statistics were performed with one-way ANOVA test. Statistical software R^64^ was used for statistical computing and graphics based on log10(AUC) values. Heat maps were plotted with pheatmap^65^. The pvclust^66^ was used for assessing the uncertainty in hierarchical cluster analysis. For each cluster in hierarchical clustering, probability values (p-values) are calculated via multiscale bootstrap resampling and raging between 0 and 1, which indicates how strong the cluster is supported by data. Two types of p-values are available: approximately unbiased (AU) p-value and bootstrap probability (BP) value. The AU p-value, which is calculated by multiscale bootstrap resampling, is superior approximation to unbiased p-value over BP value computed by normal bootstrap resampling. With pvclust hierarchical cluster analysis is performed via function hclust and automatically computes p-values for all clusters contained in the clustering of original data.

## Results

Using a rapid, semi-automated, lysis profile-based workflow analysis (Fig. 1), we assessed the impact of the antibiotics, colistin or ciprofloxacin, on *P. aeruginosa* phage PEV2 infection activities. We were then able to infer phage population growth and bactericidal activity through endpoint plating of these cultures for PFUs and CFUs. Results were then evaluated separately via one-step growth experiments. See Appendix A for discussion of colistin as well as ciprofloxacin MIC, clinical dosages, and pharmacology. See also Appendix B for detailed description of lysis profile analysis.

**Figure 1.**
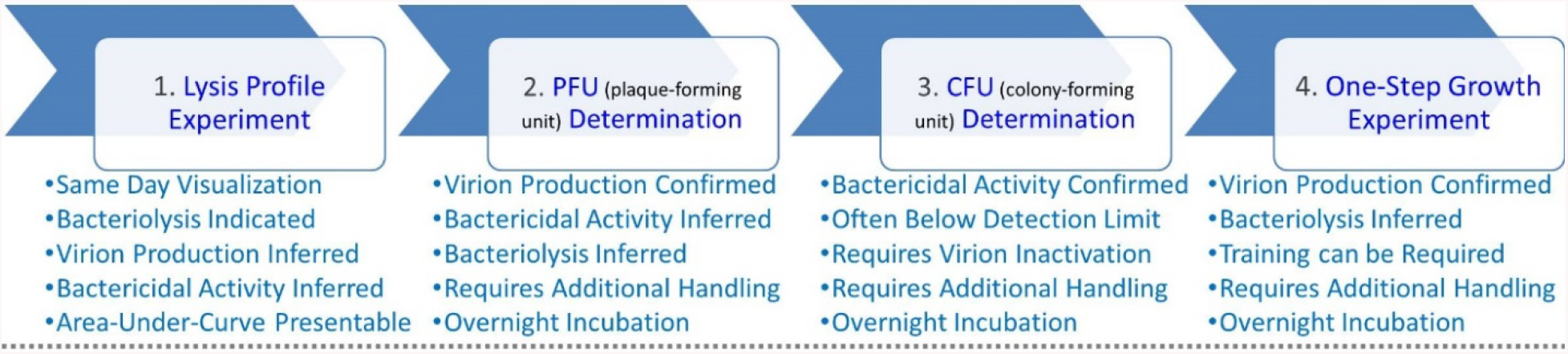
Workflow schematic. **(1.)** Lysis profile experiments can be performed employing a variety of phage multiplicities of infection (MOIs), antibiotic concentrations (MICs), and, in principle, also different phage types, antibiotic types, host variants, and conditions. Following individual lysis profile experiments, then PFU counts **(2.)** and CFU counts **(3.)** can be determined to confirm virion production and bactericidal activities, with these latter analyses requiring additional plating as well as overnight incubation. **(4.)** Phage infection activities can then be assessed independently using one-step growth experiments.

### Colistin: high levels of phage PEV2 antagonism

Colistin (polymyxin class) is a rapid acting bactericidal antibiotic that disrupts the outer bacterial membrane, leading to leakage of cell contents and death^67^. This action is achieved partially by colistin displacing the calcium and magnesium bridges that stabilize LPS^68–72^. In our study, phage PEV2 lysis profiles in the presence of colistin indicated only limited but nevertheless still clearly present phage-associated bacteriolytic activity, as seen with the MOI 5 and 10 curves at 1× MIC colistin (Fig. 2C and Fig. S1C, and see also appendix B). No phage-associated bacteriolytic activity appears to occur, however, at higher colistin concentrations (Fig. 2D and E, see also Fig. S1 and Fig. S2). The MOI 0.1 and 1 curves at 1× MIC colistin showed little or no difference relative to no phage addition, suggesting little virion production activity in addition to the low bacteriolytic activity (both as seen in Fig. 2C). These results are summarized in Figs. 2F and 2G in terms of areas under the curves (AUCs) and as heat maps based on log10(AUC) calculations (Fig. S5A). Phage PEV2 (family *Podoviridae*) utilizes an LPS surface receptor. This could be a contributor to the observed antagonism of this phage’s infection activity by colistin as LPS is directly disrupted by that antibiotic. Similar antagonism was observed, however, with the infection activity of other *P. aeruginosa* short-tailed phages from the *Autographiviridae* family, that utilize type IV pili as a surface receptor (supplementary Table 1 and Figs. S1 and S2). These results are suggestive of a general trend of phage infection activities antagonism by colistin, though are not indicative of the cause of this antagonism.

**Figure 2.**
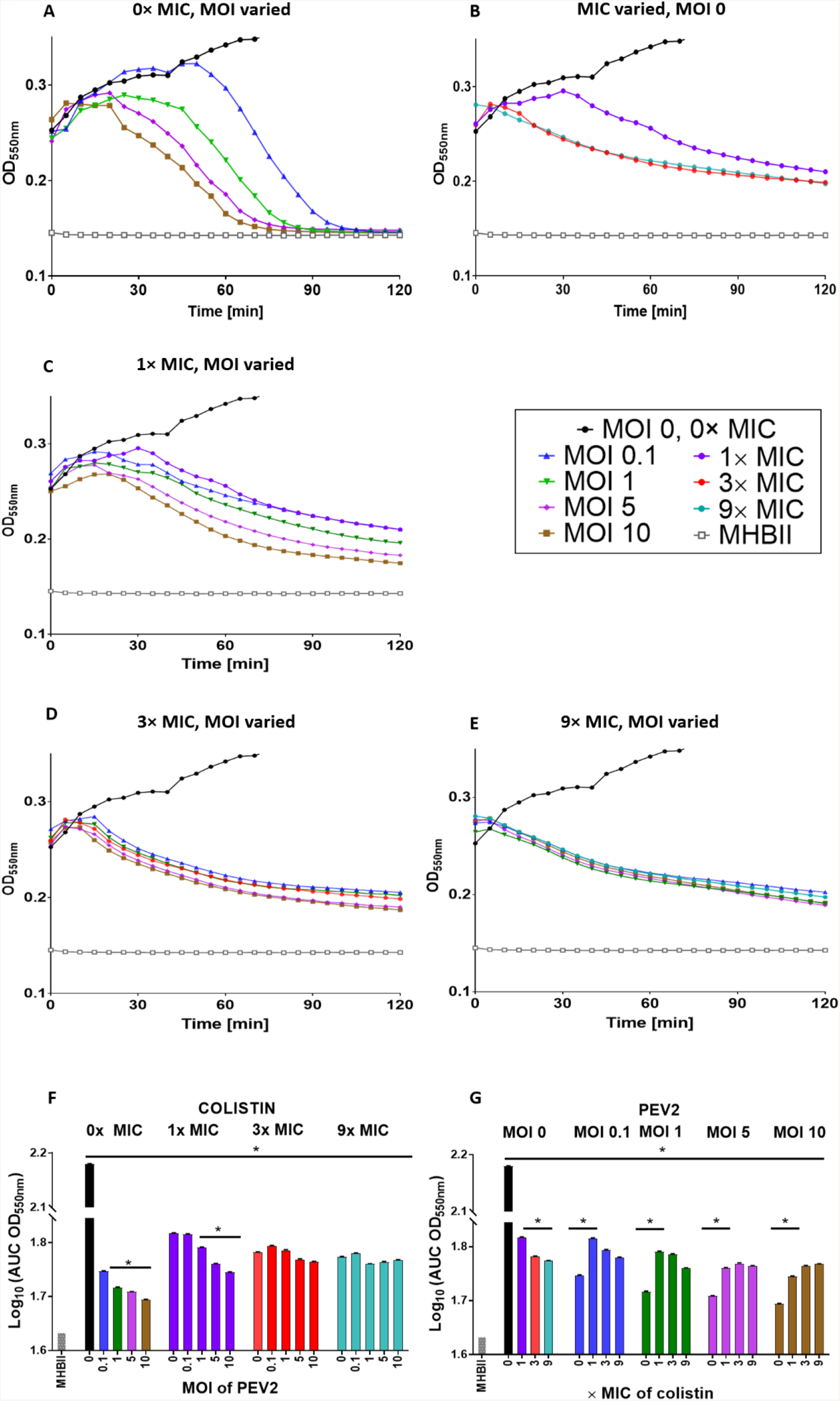
Colistin strongly antagonizes phage PEV2 infection of *P. aeruginosa*. **(A)** Phage PEV2 behavior in the absence of colistin and impact on *P. aeruginosa* PAO1 cultures. **(B)** Impact of different concentrations of colistin on *P. aeruginosa* PAO1 culture without phage. **(C-E)** Impacts of 1×, 3×, and 9× MIC concentrations of colistin, respectively, at various MOIs of PEV2 phage. **(F-G)** Area under the curve analysis of PEV2 infection with colistin. Data are derived from lysis profiles equivalent to those presented in A-E. The smaller the area under a lysis profile curve then the greater the reductions in culture turbidity over time, such as mediated by phage infection. A single representative experiment is shown. A key is provided in the black frame. Error bars represent standard deviation between samples. See Appendix A for description of the explicit colistin concentrations used and Appendix B for interpretation of lysis profile results.

### Ciprofloxacin: low levels of phage PEV2 antagonism

Ciprofloxacin, a fluoroquinolone class antibiotic, binds to DNA gyrase (atypical topoisomerase II) or topoisomerase IV in the presence of DNA^73^, prevents unlinking or decatenating of DNA following DNA replication while also negatively impacting positive supercoil relaxation and torsional stress relief ahead of transcription and replication complexes^74,75^. Representative lysis profile experiments illustrating the impact of ciprofloxacin on phage PEV2 infection are presented in Fig. 3. Contrasting with the colistin results (Fig. 2), negative impacts on lysis-profile kinetics are minimal at 1× and 3× MIC ciprofloxacin concentrations. At 9× MIC, which is slightly above the maximum serum concentration of 500 mg ciprofloxacin dose in clinic (Appendix A), bacteriolytic activity also is relatively unchanged, as seen at MOIs of 1, 5, and 10 (Fig. 3E). With MOI 0.1 and 9× MIC, however, culture-wide lysis is delayed (Fig. 3F), perhaps suggesting reduced burst sizes at this ciprofloxacin concentration. See Appendix B for discussion of the basis of these inferences.

**Figure 3.**
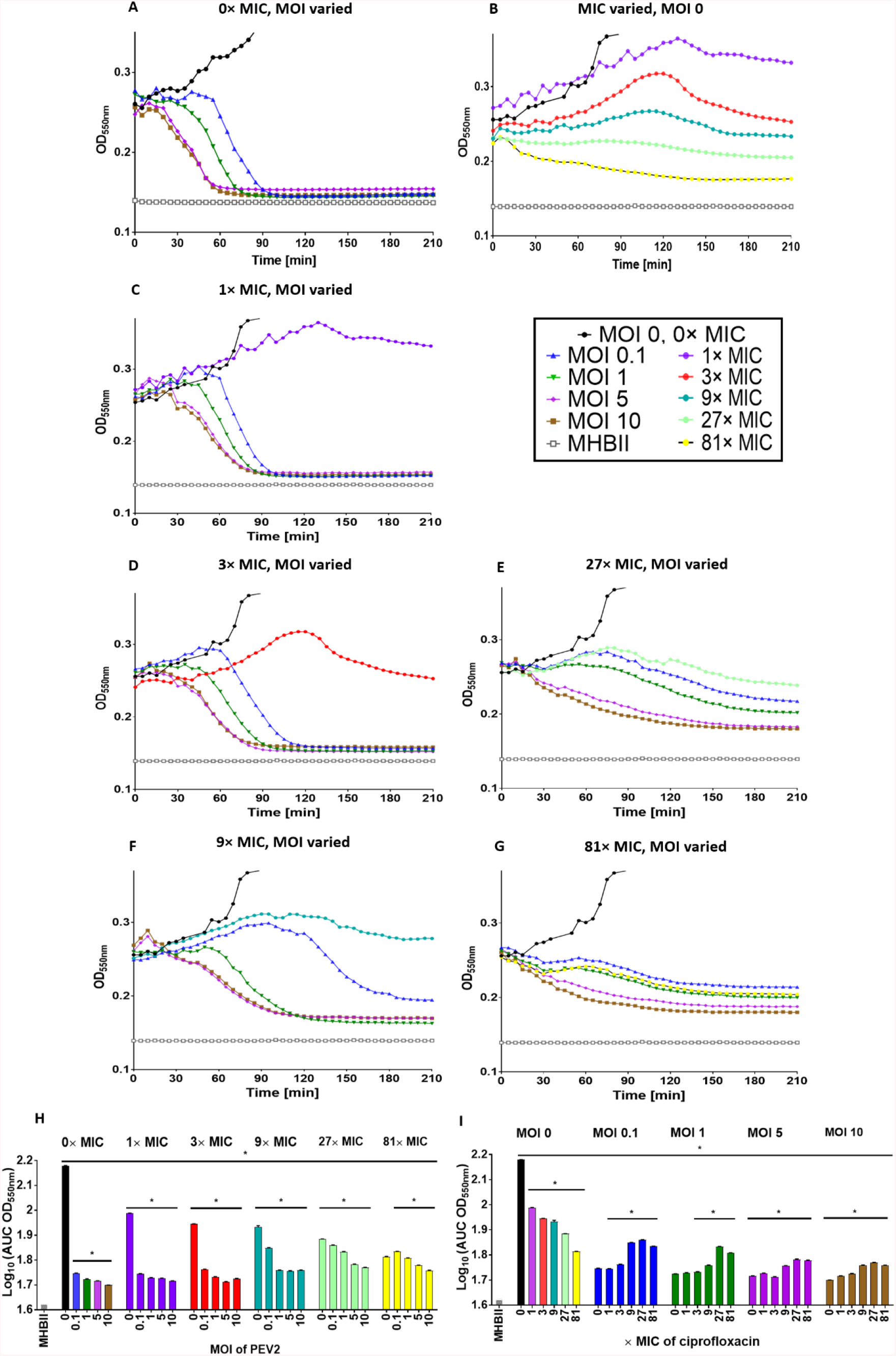
Ciprofloxacin low antagonism of phage PEV2 infection of *P. aeruginosa*. **(A)** Phage PEV2 behavior in the absence of ciprofloxacin and impact on *P. aeruginosa* PAO1 cultures. **(B)** Impact of different concentrations of ciprofloxacin on *P. aeruginosa* PAO1 cultures without phage. **(C-G)** Impacts of 1×, 3×, 9×, 27× and 81× MIC concentrations of ciprofloxacin, respectively, at various PEV2 MOIs. **(H-I)** Area under the curve analysis of PEV2 infection with ciprofloxacin. Data are derived from an equivalent to those presented in A-G. The smaller the area under a lysis profile curve then the greater reductions in culture turbidity over time, such as mediated by phage infection. A single representative experiment is shown. A key is provided in the black frame. Error bars represent standard deviation between samples. See Appendix A for explicit ciprofloxacin concentrations used and Appendix B for interpretation of lysis profile results.

At 27× MIC (8.24 μg/ml), which is above maximum serum concentration of the highest ciprofloxacin clinical dose (1000 mg, 5.4 μg/ml maximum serum concentration, see Appendix A), some phage bacteriolytic activity might still persist, and perhaps even virion production (Fig. 3F and G). Even at 81× MIC (24.72 μg/ml), almost five times maximum serum concentration of 1000 mg dose, lysis profiles suggest that it is possible that phage PEV2 bacteriolytic activity could still be present (MOIs of 5 and 10). We do not have high confidence in those conclusions as based on these lysis profile experiments alone, however. No obvious differences are seen between MOIs of 0 (i.e., no phage), 0.1, and 1 at 81× MIC ciprofloxacin concentration (Fig. 3G) suggesting little retention of virion production at such high ciprofloxacin concentrations even if phage-induced bacteriolytic activity has not been completely lost. See Figs. 4H and 4I for summary of ciprofloxacin results in terms of AUCs and Fig. S5B for heat map based on log10(AUC) calculations. We also found no difference in any of these results when phage addition was delayed by 30 min after ciprofloxacin addition (data not presented).

**Figure 4.**
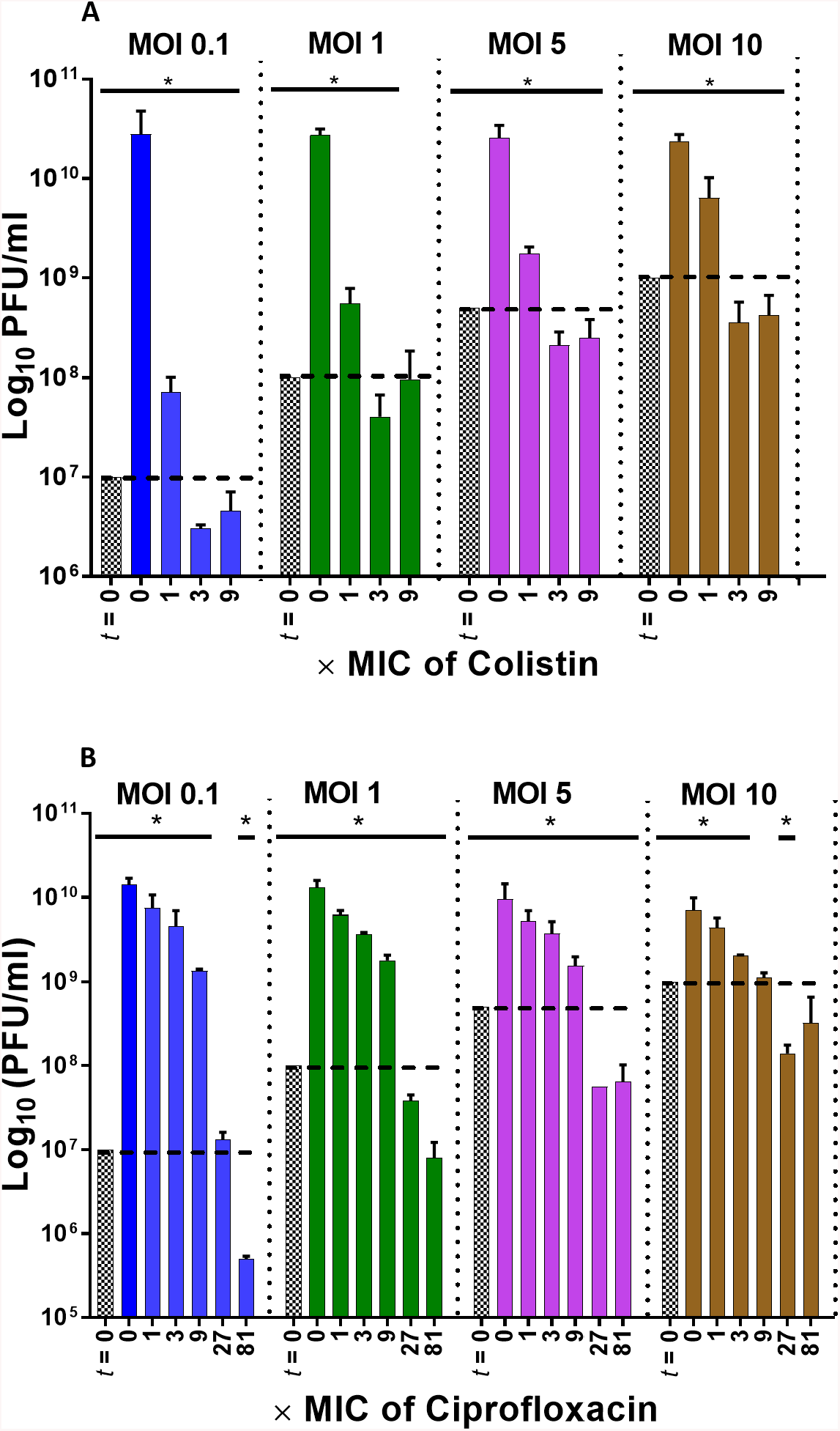
Corroboration of high antagonism of colistin and lower antagonism of ciprofloxacin on phage production. Following lysis-profile experiments, counts of phage plaque-forming units (PFUs) were determined with and without antibiotic treatments (different MICs, see bottoms of graphs) and with different starting MOIs (top of graphs). PFUs above the dashed lines indicate more phages present than at the start of experiments. Patterned bars (*t* = 0) represent initial phage titer applied at the beginning of experiments and are compared with 5h time points (post lysis profile determinations). The detection limits were 10^2^ per ml. [*] *p*<0.05, ANOVA when compared to initial (input) titer (*t* =0).

### Plaque formation support the lysis profile result

In conjunction with lysis profile experiments, it is possible to directly assess endpoint phage (PFU) viable counts. Increases in numbers of PFUs over the course of lysis-profile experiments are indicative of virion production. While virion production is expected during infections of bacteria by strictly lytic phages, it is not a given that this will occur in the presence of antibiotic.

Consistent with the lysis profile results (Fig. 2), endpoint PFU counts are substantially decreased at 1× MIC colistin in comparison to the no-antibiotic control (over two-log reductions for MOI 0.1, *p*<0.05), and reduced below input PFUs at ≥3× MIC (*p*<0.005, Fig. 4A). Also consistent with the lysis profile results, and contrasting the colistin results, ciprofloxacin at 1× and 3× MIC only minimally decreased phage PEV2 PFUs at all MOIs relative to the no-ciprofloxacin control (*p*< 0.05, Fig. 4B). Ciprofloxacin concentrations of 9× MIC, however, reduced PEV2 PFU counts by ~1 log at all MOIs tested (*p*<0.05) relative to the no-ciprofloxacin control and these levels were not significantly different from input PFUs with MOI 10 (Fig. 4B). At ciprofloxacin concentrations of 27× and 81× MIC – which are above maximum serum concentration of the highest ciprofloxacin dose in clinic (Appendix A) – drops in phage titers were observed in comparison to input concentrations of phage PEV2 for MOIs 1 through 10 (*p*<0.05) but with no change for 27× MIC and MOI 0.1.

### Bacteria elimination by phages in the presence of antibiotic

Decreases in numbers of CFUs relative to the start of lysis-profile experiments are directly indicative of treatment bactericidal activities. Phage PEV2 reduced CFU counts by at least 99.999% (4 log) in all single (phage-only) and mixed treatments (phage plus antibiotic; *p*< 0.005) over 5h (Fig. 5). Colistin treatment alone at 1× MIC reduced CFUs more than one log in comparison to all phage-only treatments (*p*<0.005), i.e., about 5 logs. Combination treatments of colistin (all MICs) and phage PEV2 (all MOIs) as well as colistin alone, at ≥3× MIC, were the most successful at reducing CFU counts, by a total of at least 6 logs. There was no significant difference between all phage treatments alone (MOIs 0.1 through 10) and ciprofloxacin treatment alone at 1× MIC. At 3× MIC, ciprofloxacin alone reduced CFU numbers to a greater extent (over 1 log more, *p*<0.05) than phage-alone treatments. As in case of colistin, co-treatment with the use of phages (all MOIs) and ciprofloxacin (all MICs) were the most successful at reducing CFU counts (≥ 6 logs).

**Figure 5.**
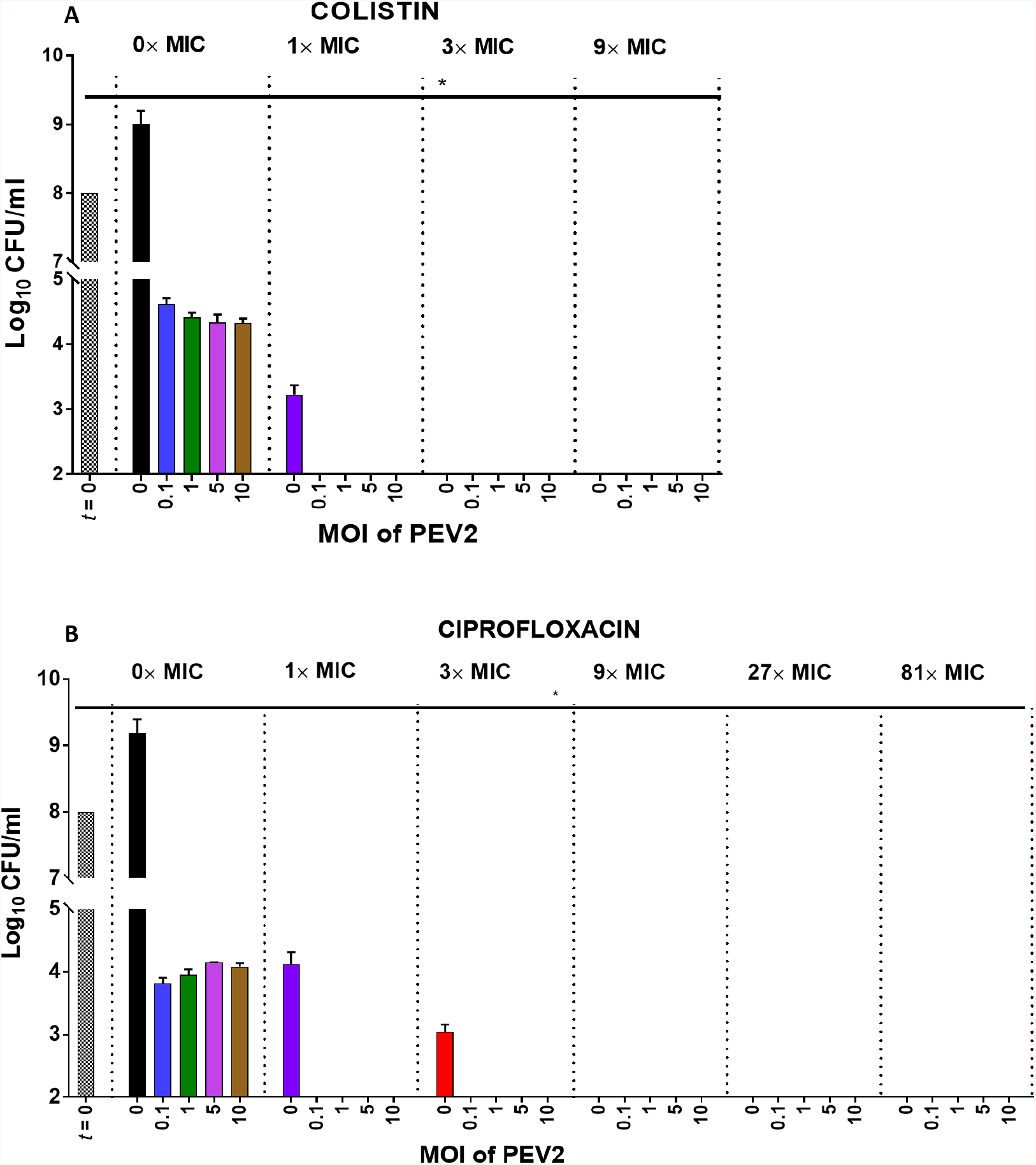
Higher impact of phage-antibiotic combination treatments. Following lysis-profile experiments, counts of bacterial colony-forming units (CFUs) were determined with and without phage treatments (different MOIs, see bottoms of graphs), antibiotic treatments (different MICs, see tops of graphs), and combined treatments. CFUs bars that are below the black bar (mock treatment, no phages and no antibiotics) indicate reductions in bacterial numbers. To prevent bactericidal phage adsorption during enumeration, CFU analyses were performed in the presence of free phage-inactivating virucide consisting of 7.5% black tea infusion & 0.53% FeSO_4_^58^. [*] *p*< 0.005, ANOVA. All the missing bars represent less than 10^2^ CFUs/ml, which is the detection limit of this assay.

### One-step growth experiments corroborate lysis profile results

For ~80 years the standard method of assessing phage infection characteristics, particularly determining phage burst size and latent period, has been using one-step growth experiments^62,76,77^. These differ from lysis profiles both in terms of the potential for lysis-released virions to adsorb already phage infected bacteria (high potential with lysis profiles vs. low potential with one-step growth experiments) and the means of enumeration (culture turbidity for lysis profiles vs. via plaque counts for one-step growth experiments). Importantly, one-step growth analysis is also a more complicated and time-consuming assay (weeks for optimization to specific phages, bacteria, and conditions) in comparison to the presented lysis-profile analyses (hours, i.e., Figs. 2 and 3).

To test the accuracy of lysis profiles as predictors of antibiotic antagonistic impacts on phage PEV2 infection activity, we also employed one-step growth experiments. At 1× and 3× MIC, colistin did not have a significant impact on the phage PEV2 latent period (Fig. 6D). At higher colistin concentrations (>3× MIC), lawn growth during PFU assessment was inhibited, presumably by antibiotic carry over and therefore one-step growth assessment could not be performed (this was also seen at 81× MIC ciprofloxacin but was not an issue post lysis-profile experiments, i.e., Fig. 4, due to greater degrees of dilution prior to plating). Contrasting the consistency of latent periods seen in the presence of colistin, burst sizes were substantially reduced (*p*<0.005) at 1× and 3× MIC colistin, to 9.6 ± 0.57 (92.9% of reduction) and 2.5 ± 0.2 (98.1% of reduction), respectively (Fig. 6D and E).

**Figure 6.**
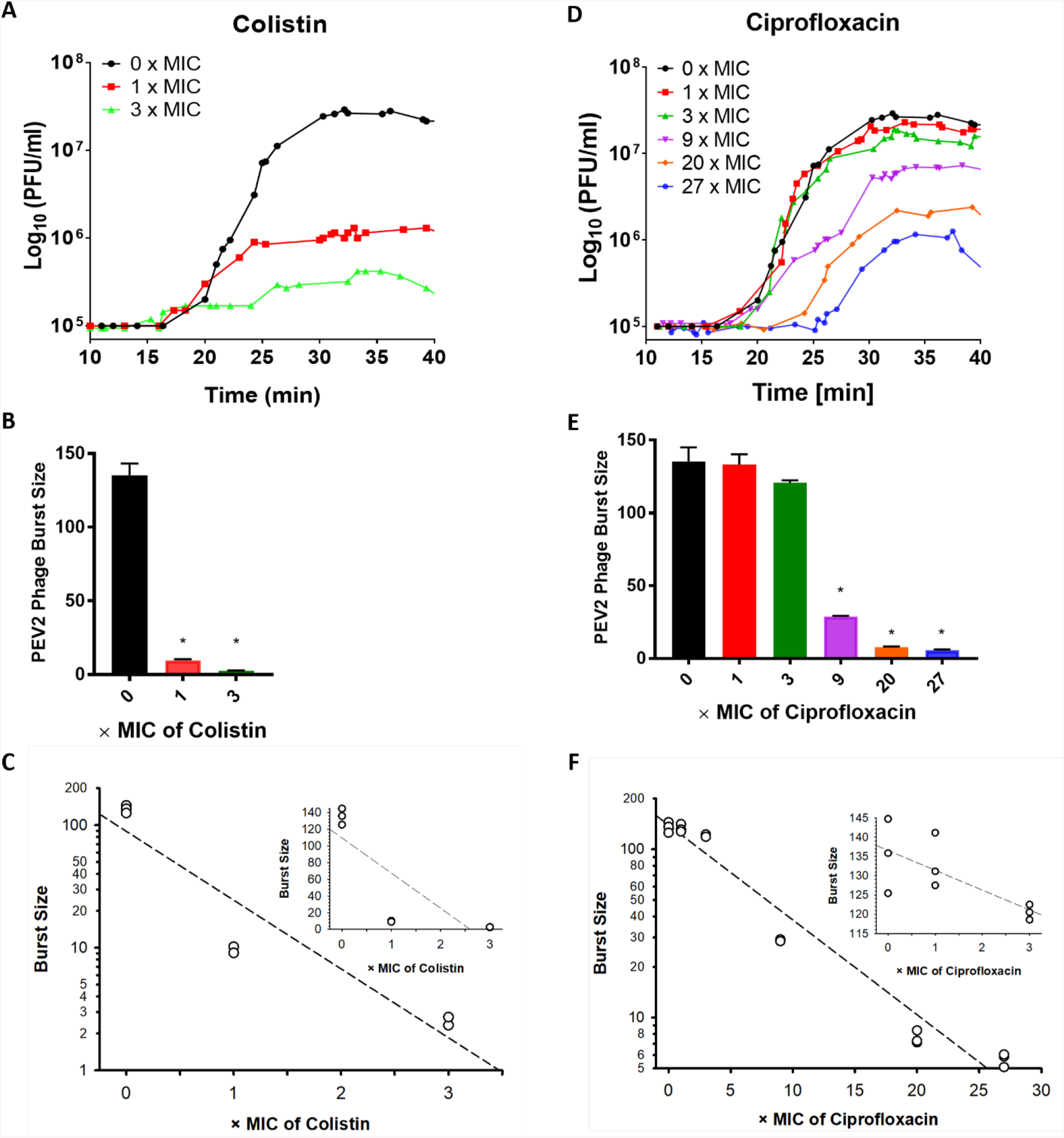
PEV2 one-step infection cycle characteristics in the absence or presence of antibiotic. One-step growth curve of PEV2 infection (MOI 0.05) in the absence or presence of colistin **(A)** or ciprofloxacin **(D)**; PEV2 phage burst size in the absence or presence of different colistin **(B)** or ciprofloxacin **(E)** concentrations. Error bars represent standard deviations between three experiments (biological replicates); relationship between burst size and colistin (**C**, *m*= −0.561, *r*= 0.938) or ciprofloxacin (**F**, *m*= −0.056, *r*= 0.982) concentrations. Inserts show burst sizes using non-log scales at colistin or ciprofloxacin first three concentrations (0×, 1×, and 3× MIC). The dotted line represents a trend line. (*) *p* < 0.005, ANOVA, (*r*) linear regression, (*m*) slope. See Appendix A for detailed antibiotics concentrations.

As seen also with colistin, when *P. aeruginosa* PAO1 was incubated for 15 min with ciprofloxacin at 1×, 3× or 9× MIC concentrations and then infected by phage PEV2, the latent period did not change. Burst sizes, however, were 133.2 ± 4.1, 120.5 ± 1.1, 28.8 ± 0.3 phage PFUs per infected bacterial cell (mean values from 3 separate experiments), corresponding to 1.5%, 10.9% and 78.7% reductions in burst size, respectively (Fig. 6B and C). When ciprofloxacin concentrations were increased to 20× and 27x MIC (which are above the highest clinical serum concentrations; see Appendix A), the PEV2 latent period was postponed by 3 and 7 min, respectively, and burst size dropped to 7.6 ± 0.4 and 5.7 ± 0.3 phages/infected bacterium, the latter corresponding to 94.4 % and 95.8 % reductions in burst size. Figs. 6C and 6F present log-linear plots of declines in burst size of phage PEV2 as a function of ciprofloxacin or colistin concentrations.

These one-step growth results are consistent with the lysis profile results presented in Figs. 2–5, indicating for example that ciprofloxacin does not excessively interfere with phage infection activities until relatively high antibiotic concentrations are present (>>9× MIC) while colistin does substantially interfere even at relatively low concentrations (1× MIC). Both bacteriolysis and virion production seem to be present in lysis profiles at 27× MIC of ciprofloxacin and 1× as well as 3× MIC of colistin, though nearing the limits of detection, which would appear to be consistent with the observed retention in one-step growth experiments of low but still present burst sizes. Similarly, the low burst size seen at 27× MIC of ciprofloxacin and 3× MIC of colistin seems to be consistent with the lack of detection of increases in PFUs following lysis profiles in the presence of these concentrations of antibiotic (Fig. 4).

## Discussion

In combination therapies, two or more agents possessing different mechanisms of action can enhance each other’s activities^78–80^, broaden activity spectra^78,81–83^, reduce the potential for resistance evolution^84–91^, or improve penetration into biofilms^92^. One common aspect of phage therapy treatments is combining phages with antibiotics^38,92–102^. Here we have assessed the potential of different concentrations of two different classes of antibiotics, colistin and ciprofloxacin, to interfere with infection activities of *P. aeruginosa* phage PEV2 and other tested phages at bacterial growth-inhibiting antibiotic concentrations (≥MIC).

### High antagonistic impact of colistin

David *et al.*^103^ showed that colistin at 1× MIC (1 μg/ml) inhibited phage production during infection of *Mycobacterium aurum*. Jansen *et al.*^100^ on the other hand reported phage-mediated reductions of *Acinetobacter baumannii* culture growth in the presence of colistin, as determined by optical density and even staring with low MOIs. They also observed little reduction in culture growth in the presence of colistin alone, however, which could suggest phage infection activity in the presence of either colistin-resistant or colistin-tolerant bacteria, rather than in the presence of fully colistin-sensitive bacteria.

Using a cocktail of two phages, Chaudhry *et al.*^99^ observed substantial phage population growth as well as phage-associated reductions in *P. aeruginosa* (PA14 vs. PAO1 as used here) in the presence of 8× MIC (20 μg/ml) colistin. In these experiments, however, phages were targeting bacterial biofilms rather planktonic bacteria, and biofilms tend to be more tolerant of colistin than bacteria found in broth^104^. Perhaps consistently, MICs were not determined in this study using biofilm-grown bacteria and neither 1× nor 8× MIC colistin alone had much of a negative impact on the treated biofilms. With similar caveats, Danis-Wlodarczyk *et al.*^59^ observed that application of the giant *phiKZvirus* KTN4 together with 100 μM (116 μg/ml) of colistin substantially reduced *P. aeruginosa* PAO1 72h old biofilm, while colistin alone did not have an impact and phage-alone treatment eradicated biofilm to a lesser extent than the combination treatment.

Overall, then, evidence of substantial phage replication while infecting bacteria that are truly colistin-inhibited in their growth is not robust. Done under conditions resembling those employed for MIC determinations, we nonetheless have equivalently found that colistin is strongly antagonistic toward phage PEV2 bacteriolytic and virion production activities, e.g., with greater than 90% reduction in burst size at 1× MIC.

### Low antagonism of ciprofloxacin

A handful of studies have evaluated the impact of ≥1× MIC ciprofloxacin concentrations on the *in vitro* infection activities of various phages. These were generally done over many rounds of phage infection, however. In addition, with few exceptions, these examinations have been done under conditions not equivalent to those used for MIC determinations, i.e., as biofilms are expected to display greater tolerance to ciprofloxacin activity than broth-growing bacteria^104^. Thus, and just as with colistin (above), ciprofloxacin concentrations that are found to be inhibitory of bacterial growth in broth may not be inhibitory to bacterial metabolic activities within biofilms.

Verma *et al.*^105^ saw no change in phage impact on *Klebsiella pneumoniae* biofilms with and without ciprofloxacin (10 μg/ml, >>1× MIC), suggesting a retention of phage infection activities in the presence of this antibiotic. The phage used appeared to produce an anti-biofilm depolymerase enzyme, however, which could have reduced biofilm presence without corresponding phage infection activity. Working with *P. aeruginosa* PA14 biofilms, Chaudhry *et al.*^99^ found that 8× MIC (6.4 μg/ml) ciprofloxacin blocked the production of new virions while 1× MIC (0.8 μg/ml) did not, though again MIC was determined using planktonic rather than biofilm cultures. Perhaps consistent with their observations of phage infection activity while in the presence of 1× but not 8× MIC ciprofloxacin, while 1× MIC ciprofloxacin alone had little impact on biofilm presence, 8× MIC ciprofloxacin acting alone had a substantial impact. Also working with ciprofloxacin and biofilms, in this case of *Staphylococcus aureus*, Dickey *et al.*^106^ found that 2× MIC (0.375 μg/ml) ciprofloxacin reduced phage population growth while 10× MIC (1.875 μg/ml) eliminated phage population growth altogether. As was also the case with the Chaudhry *et al.* observations, Dickey *et al*. found that 2× MIC ciprofloxacin acting alone had no impact on biofilm presence whereas 10× MIC ciprofloxacin had a substantial negative impact. Jansen *et al.*^100^ working with an *A. baumannii* phage and ciprofloxacin had results that were similar to their efforts with colistin, i.e., retention of phage antibacterial activity, but also with caveats equivalent to those discussed above for colistin.

Luscher *et al.*^107^ treated 6 h *P. aeruginosa* PAO1 infections of Calu-3 epithelial cell lines in minimal essential medium with dual phage treatment in the presence of 16× MIC ciprofloxacin (4.0 μg/ml, which roughly corresponds to maximum serum concentration of 750 mg dose of ciprofloxacin in clinic, 4.3 μg/ml, and corresponds with 14× MIC in our study). After 72h post infection they observed substantial bacteria killing and multiple log increases in PFUs. Only a few PAO1 mutants were isolated following treatments, with all of these mutants ciprofloxacin sensitive and all resistant to to some or all phages. The time of treatment and these resistance results highly suggest that at the start of treatments, which was six hours following epithelial cell infection, the PAO1 cells were not in a planktonic state anymore but instead had formed aggregates with biofilm-like properties and, as with the studies discussed above therefore, with possibly increased tolerance to ciprofloxacin. Additionally, ciprofloxacin can positively diffuse and is taken up actively by Calu-3 epithelial cells^108,109^, lowering the effective ciprofloxacin concentration.

Valério *et al.*^110^, contrasting the retention of phage infection activities while infecting biofilms in the presence of ciprofloxacin as discussed above, appear to have seen an absence of phage activity when non-biofilm *Escherichia coli* was treated with 1× MIC (0.5 μg/ml) ciprofloxacin. Also with *E. coli*, Lopes *et al.*^111^ observed less phage population growth in tryptic soy broth in the presence of 1× MIC (0.25 mg/mL) ciprofloxacin than without, and no increases in phage numbers were seen at 2× MIC (0.5 mg/ml). Additional but potentially less informative studies exploring the impact of ≥1× MIC ciprofloxacin on phage infection activity including in animals are reviewed by Abedon^94^ along with evidence that nalidixic acid, which displays a similar antibacterial mechanism as ciprofloxacin, is inhibitory to phage infection activity.

Exceptional among studies exploring the impact of ciprofloxacin on phage infection activities is that of Oechslin *et al.*^112^. Oechslin *et al.*^112^ found synergistic impacts of ciprofloxacin and a phage cocktail on experimental *P. aeruginosa* endocarditis in rats. This is notable since ciprofloxacin alone displayed substantial antibacterial activities in this *in vivo* system and thus presumably exceeded MIC there. Using a tryptic soy broth model, Oechslin *et al.*^112^ also found that numbers of *P. aeruginosa* cells were substantially reduced (~6 logs) after 6h of exposure to co-treatments of a phage cocktail (MOI 1, 10^8^ PFUs/mL) and ciprofloxacin (2.5× MIC, 0.475 μg/ml) as well as by phage-only treatment, versus only an inhibition of bacterial growth when treated by ciprofloxacin alone. This result is suggestive of phage virion production in both the presence and absence of 2.5× MIC ciprofloxacin since the starting MOI of 1 at best should kill only 63% of the bacteria present without considerable phage replication^113^.

Here, we have found that ciprofloxacin has a low antagonistic impact on phage PEV2 infection activity, including on virion production, even at high antibiotic concentrations. This infection activity of phage PEV2 as well as that of other tested phages is retained not only under conditions equivalent to standard broth-based antibiotic MIC determinations, i.e., within planktonic MBH-grown cultures, but at clinically relevant ciprofloxacin concentrations as well (Appendix A). This was confirmed by lysis profile and one-step growth experiments (Figs. 3 and 6), where phage PEV2 maintained over 20% of its burst size even at 9× MIC ciprofloxacin. At ≥20× MIC, which exceeds maximum serum concentrations of ciprofloxacin following the highest FDA recommended dose of this drug (Appendix A), we found that phage bacteriolytic as well as virion production activities were still retained, albeit at lower levels. These findings suggest that ciprofloxacin at high doses may not excessively interfere with phage therapies even when targeted *P. aeruginosa* is substantially affected by this antibiotic, and point to a utility to more rigorous assessments of antibiotic antagonistic activities on phages during phage therapy development.

## Conclusions

Unlike most other studies that have found some compatibility between ciprofloxacin and phage treatments, here we measured antagonism under conditions that are equivalent to determinations of MICs, including use of planktonic bacteria and MHB II media. We found that ciprofloxacin could be well suited for antibiotic-phage therapy co-treatment vs. colistin as based on phage PEV2 displaying substantial bactericidal, bacteriolytic, and phage production activities even at clinically high levels ciprofloxacin. Methodologically, we advance lysis profiles as a rapid and inexpensive broth-based assay for screening for antibiotic antagonistic impacts on phage infection activities given consistency between observations made using lysis profiles and standard one-step growth assays.

## Acknowledgements

Funding was provided by the Ohio State University President’s Postdoctoral Scholars Program and Cystic Fibrosis Foundation C3 training award (to K.D.W.), a Gordon and Betty Moore Foundation Investigator Award (#3790, to M.B.S.), and PHS funding R01AI34895 and R01AI43916 (D.J.W.).

## Author Contribution

K.D.W. performed all experiments with help of A.C. A.Ch. and M.G. helped with optimization of one-step growth experiments. K.D.W. and S.T.A. designed experiments and analyzed results. K.D.W., S.T.A., D.J.W., and M.B.S. wrote the manuscript.

## APPENDIX A Antibiotic minimum inhibitory concentrations, dosing, and pharmacology

## Colistin

Colistin can be administrated parenterally, intravenously, or via inhalation. Currently there are no clear dosing guidelines, pharmacokinetics, or pharmacodynamics provided by the FDA. In Table A1 we present recommended dosing and target serum concentrations based on the most comprehensive study of colistin treatment of ill patients and recently published International Consensus Guidelines for the Optimal Use of Polymyxins^114^.

Since there are no clear guidelines for colistin treatment, in our study we performed first a standard broth-based MIC analysis with *P. aeruginosa* PAO1 reference strain (1 × MIC = 39.06 μM = 68.75 μg/ml) and we employed the following concentrations in further experiments: 1× MIC = 39.06 μM = 68.75 μg/ml, 3× MIC = 117.18 μM= 206.25 μg/ml, 9× MIC = 351.54 μM = 618.75 μg/ml, 20 × MIC = 18.2 μM = 6.1 μg/ml, 27 × MIC = 24.70 μM = 8.24 μg/ml, and 81 × MIC = 74.10 μM = 24.72 μg/ml.

## Ciprofloxacin

In the clinic, ciprofloxacin is administrated in 2 forms, film-coated tablets, available in 100 mg, 250 mg, 500 mg and 750 mg strengths, or oral suspension, available in 5% (5 g ciprofloxacin in 100 mL) and 10% (10 g ciprofloxacin in 100 mL) strengths^115^. Ciprofloxacin maximum serum concentrations and AUC in correlation to dose are presented in Table A1, based on FDA directions for ciprofloxacin dosing and pharmacology^115^.

For this study, a standard broth-based MIC analysis was performed, which indicated that ciprofloxacin inhibits macroscopically growth of sensitive *P. aeruginosa* PAO1 Krylov strain at 305.2 pg/ml (0.91 μM). Based on these results and FDA ciprofloxacin dose recommendations, we established concentrations span for further experiments as follows: 1 × MIC = 0.91 μM = 303.2 ng/ml, 3 × MIC = 2.74 μM = 909.5 ng/ml, 9 × MIC = 8.23 μM = 2.73 μg/ml, 20 × MIC = 18.3 μM = 6.1 μg/ml, 27 × MIC = 24.70 μM = 8.19 μg/ml, and 81 × MIC = 74.11 μM = 24.56 μg/ml.

**Table A1.** Colistin and ciprofloxacin pharmacology.

| Antibiotic | Dose | Maximum serum concentration | Area Under Curve (AUC) |
| --- | --- | --- | --- |
| **Colistin (polymixin E)\*** | <sup>2</sup>75- 410 mg | 0.48- 9.38 μg/ml | N.D. |
| | <sup>3</sup>300–360 mg CBA for a patient with normal renal function | ∼2 μg/ml<sup>114,116,117*</sup> | ∼50 mg•hr/mL** |
| **Ciprofloxacin<sup>1</sup>** | 250 mg | 1.2 μg/ml | 4.8 μg•hr/mL |
| | 500 mg | 2.4 μg/ml | 11.6 μg•hr/mL |
| | 750 mg | 4.3 μg/ml | 20.2 μg•hr/mL |
| | 1000 mg | 5.4 μg/ml | 30.8 μg•hr/mL |

<sup>1</sup> presented based on FDA dosing analysis and recommendations<sup>115</sup>.
<sup>2</sup> Garonzik *et al.*, 2011<sup>117</sup>. Their recommends were against exceeding 300 mg “colistin base activity” per day (about 5 mg/kg/day) for either the loading or maintenance doses.
<sup>3</sup> Based on International Consensus Guidelines for the Optimal Use of the Polymyxins: Endorsed by the American College of Clinical Pharmacy (ACCP), European Society of Clinical Microbiology and Infectious Diseases (ESCMID), Infectious Diseases Society of America (IDSA), International Society for Antiinfective Pharmacology (ISAP), Society of Critical Care Medicine (SCCM), and Society of Infectious Diseases Pharmacists (SIDP) <sup>114</sup>
**CBA:** colistin base activity
**N.D.:** not determined
\* the target average steady-state total plasma colistin concentrations ( $C_{ss,avg}$ ) of ≥ 2.5mg/L may increase the risk of nephrotoxicity (controversial in the field), that complicates the clinical decision to choose the appropriate colistin dosing<sup>118,119 120</sup>
\*\* maximum tolerable exposure

## APPENDIX B Interpretation of lysis profiles

Phage infection characteristics can be inferred from lysis-profile experiments, with different aspects assessed given different starting (input) phage MOIs. These, as seen with phage PEV2 without antibiotic treatments, are summarized in Fig. B1 and below. Specifically, MOIs of 0.1, 1, 5, and 10 have been employed throughout this study, which can provide evidence of virion production, phage-induced bacterial lysis (i.e., lysis from within^51^), and phage lysis timing (latent period), respectively.

**Figure B1.**
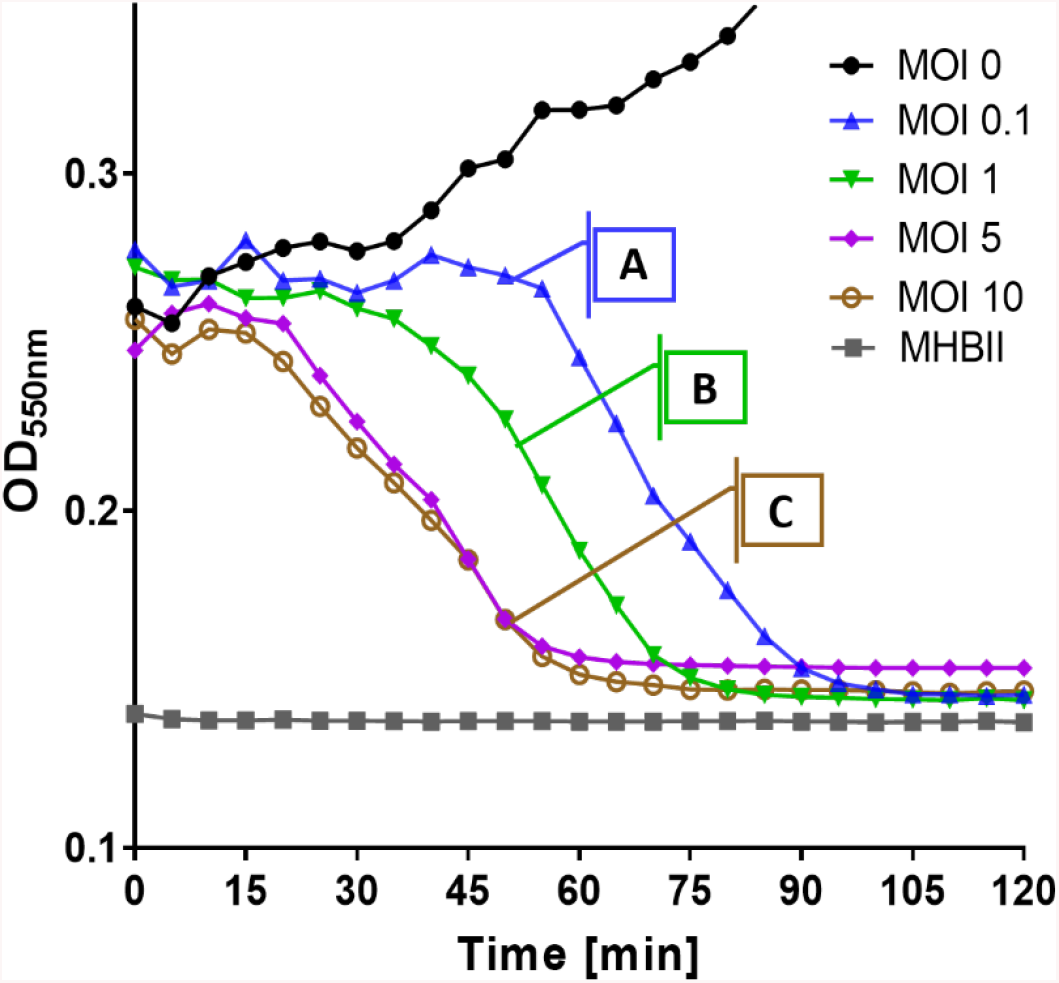
Phage PEV2 lysis profiles. **(Curve A)** Low starting MOI (= 0.1): Phages must replicate and produce new virions to give rise to culture-wide bacterial lysis, as will occur after a minimum of 2 infection cycles. **(Curve B)** Intermediate MOI (= 1): Culture-wide bacterial lysis is unlikely to be due to lysis from without (an extracellularly, enzymatically induced lysis of bacterial cells following high-multiplicity phage adsorptions), though lysis may be delayed relative to curve C due to delayed and only fractional virion adsorption. **(Curve C)** High starting MOI (5 or 10): Timing of bacterial lysis should occur after a single phage latent period, unless somewhat sooner due to lysis from without. For especially MOIs 5 or 10, phage infections need only retain bacteriolytic activity to result in robust drops in turbidity.

## Curve A. Indicating moderate or greater virion production

Starting with an input MOI of 0.1, the timing of lysis should begin after no less than two consecutive phage latent periods rather than just one (curve A, Fig. B1). This is because initially, at this MOI, only about 10% of the bacterial population can become phage infected, and it will take one latent period (the first latent period) before new virions are released. These new virions, if produced in sufficient numbers, can then adsorb and infect additional bacteria. Such ‘two-step’ growth can then result in relatively rapid culture-wide lysis, but only if phage burst sizes are sufficiently large to result in infection of the majority of bacteria present. That is, such as in excess of ~10 phages/infected bacterium so that most of the bacteria in the culture will have become phage infected after the first phage latent period. Observations of timely culture-wide reductions in turbidity at this low MOI thus can serve as an indication of phage virion production in the presence of antibiotic.

## Curve B. Indicating lysis from within

With MOI 1 (curve B, Fig. B1) we observe slower lysis than with MOIs 5 or 10 (curve C, Fig. B1). This delay could be a consequence of slow initial phage adsorption to bacteria, as with an input MOI of 1 the rate that a bacterium becomes adsorbed by at least one phage virion should be five-to ten-fold slower than with the higher input MOIs. In addition, with MOI 1 only approximately one-half [63% = 100 × (1 – e^-1^)] of bacteria will be initially infected, i.e., as due to Poissonal distributions of phage adsorption^113^. Slower culture-wide lysis at the end of phage latent periods thereby can result both because fewer bacteria are infected and bacteria are infected more slowly relative to cultures initiated with MOIs of 5 or 10. Notwithstanding such delays, culture-wide clearing for MOI 1 curves should be indicative of an occurrence of lysis from within as under these low-multiplicity conditions lysis from without (below) is unlikely.

## Curve C. Indicating timing of lysis

The timing of lysis in lysis profiles, especially given starting MOIs of somewhat greater than 1 (here MOI 5 or 10), can approximate phage latent periods. This is seen here as the initial drop in turbidity observed soon after 15 min with the curves labeled as curve C in Fig. B1. These higher input MOIs under certain circumstances, perhaps especially given antibiotic presence, might instead result from lysis from without. That is, a rapid, external digestion and then lysis of bacterial cells walls following high multiplicity virion adsorption^121^. Contrast instead lysis from within, which is phage-induced bacterial lysis as it normally occurs at the end of a phage’s latent period^122^, i.e., as with greater certainty is what is observed with given lower MOI experiments (curves A and B, Fig. B1).

