## Supplementary data for "*In vitro* analysis of colistin and ciprofloxacin antagonism of *Pseudomonas aeruginosa* phage PEV2 infection activities"

### Phage PEV2 infection characteristics

Spot test analysis employing different isogenic *P. aeruginosa* PAO1 surface mutants revealed that LPS, specifically O-specific antigen (OSA, formerly termed B-band) serves as an essential bacterial cell surface structure for successful phage PEV2 adsorption (Table S1).

**Table S1. Phage PEV2 receptor identification using isogenic *P. aeruginosa* PAO1 mutants.**

| | **Bacterial strain** | **Phenotype** | **Origin** | **Sensitivity to PEV2** |
|---|---|---|---|---|
| **Wild Type** | PAO1 Krylov | Wild type | Queen Astrid Military Hospital, Belgium, Jean-Paul Pirnay | + |
| | PAO1 (ATCC 15692) | Wild type | American Type Culture Collection, Jean-Paul Pirnay | + |
| | PAO1 | Isogenic wild type | Harvard University | + |
| **Type IV deficient** | PAO1 Pirnay | Wild type with inactive type IV pili | Queen Astrid Military Hospital, Belgium, Jean-Paul Pirnay | + |
| | PAO1Δ*pilA* | Lack of type IV pili | University of Washington, USA Matthew R. Parsek | + |
| **Flagella deficient** | PAO1Δ*fliC* | Lack of flagella | University of Calgary, Canada, Joseph Harrison | + |

| | | | | |
|---|---|---|---|---|
| **Flagella & Type IV deficient** | PAO1Δ*fliC*Δ*pilA* | Lack of flagella; lack of type IV pili | University of Calgary, Canada, Joseph Harrison | + |
| **LPS variants** | PAO1Δ*gmd* (CPA-) | Lack of *gmd* gene responsible for the biosynthesis of GDP-D-Rha, the nucleotide sugar precursor for CPA | University of Guelph, Canada, Joseph S Lam[58] | + |
| | PAO1Δ*rmlC* (OS-) | Lack of *rmlC* gene, responsible for TDP-L-Rha biosynthesis that lead to defective core OS truncated at the Rha$^{A}$ and Rha$^{B}$ residues in the two glycoforms of the OS | University of Guelph, Canada, Joseph S Lam[58] | - |
| | PAO1Δ*wbpW* (CPA-) | Lack of *wbpW* gene that encode enzymes responsible for the biosynthesis of GDP-D-Rha, the nucleotide sugar precursor for CPA | The Ohio State University, USA, Daniel Wozniak | + |
| | PAO1Δ*wzy* (OSA-) | Lack of OSA polymerization of LPS. LPS consists OS and one OSA. CPA is intact | University of Guelph, Canada, Joseph S Lam[58] | - |
| | PAO1Δ*waaL* (CPA-, OSA-) | Lack of WaaL ligating O-polymer to core-lipid A; LPS is devoid of CPA and OSA, semi rough (SR-LPS, or core-plus-one O-antigen) | University of Guelph, Canada, Joseph S Lam[58] | - |
| | PAO1Δ*algC* | Lack of *algC* required for CPA, core oligosaccharide, and alginate biosynthesis | Emory University, USA, Joanna Goldberg | - |

"+" indicates: allows clear spot formation (sensitive strain to phage treatment and presumably phage adsorption);
"-": indicates: does not allow spot formation (not sensitive strain to phage treatment and presumably phage adsorption);
CPA: LPS common polysaccharide antigen (formerly termed A-band);
GDP-D-Rha- nucleotide precursor for CPA;
L-Rha: **L**-rhamnose;
LPS: bacterial lipopolysaccharide;
OS: the LPS core oligosaccharide;
OSA: O-specific antigen (formerly termed B-band);
Rha$^{A}$ and Rha$^{B}$ : residues in the two glycoforms of the core oligosaccharide of LPS;
TDP-L-Rha: a sugar donor and core oligosaccharide of LPS

### Lysis profiles

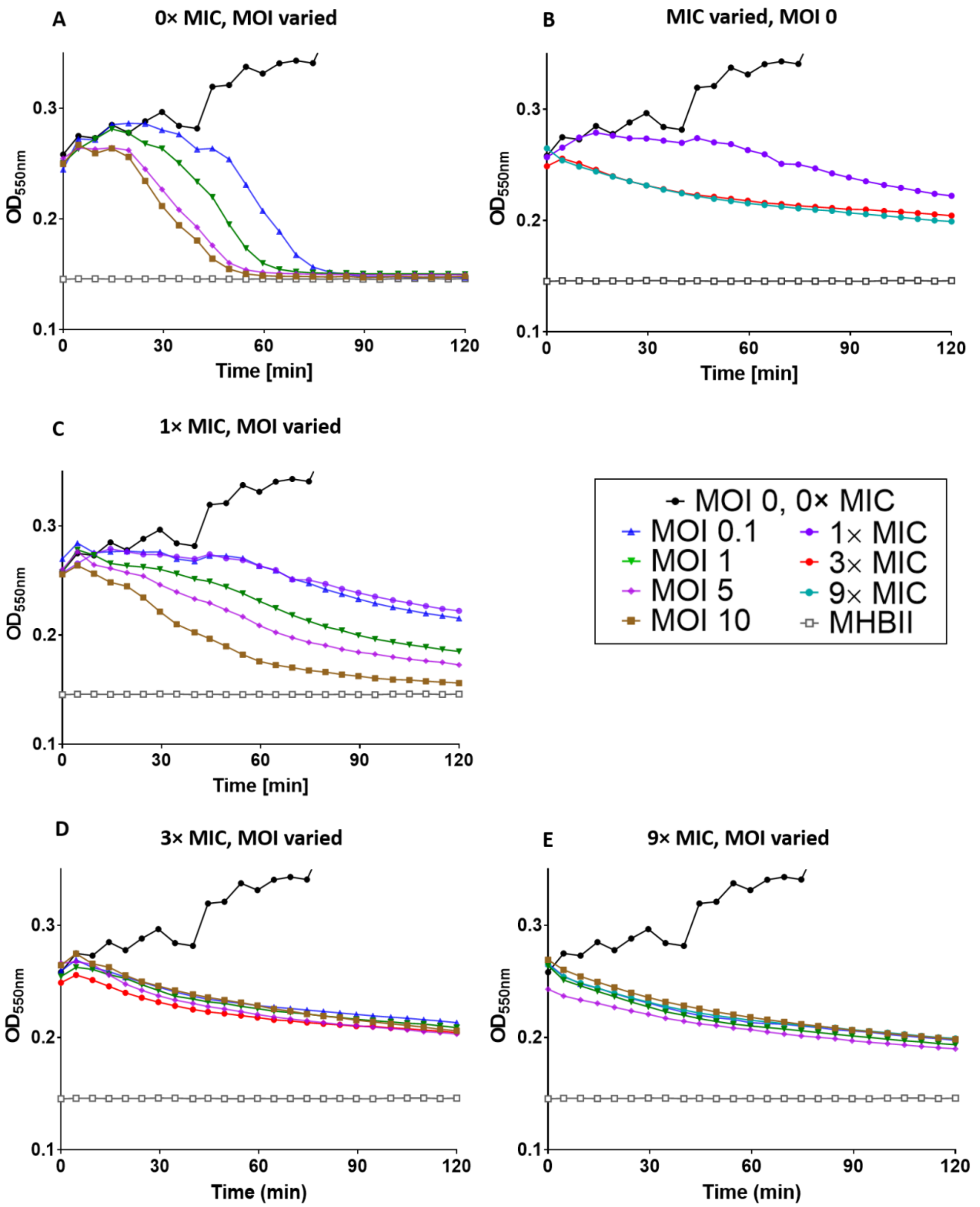


**Figure S1. High antagonism of colistin on LUZ19 infection activities. (A)** Phage behavior in the absence of antibiotic and impact on *P. aeruginosa* PAO1 cultures. **(B)** Impact of different concentrations of antibiotic on *P. aeruginosa* PAO1 culture without phage. **(C-E)** Impacts of various antibiotic concentrations (incidated as × MICs) in combination with various phage MOIs. A single representative experiment is shown. A key is provided in the black frame. See Appendix A for description of the explicit antibiotic concentrations used. LUZ19 belongs to *phiKMVvirus* and uses type IV pili as a surface receptor.

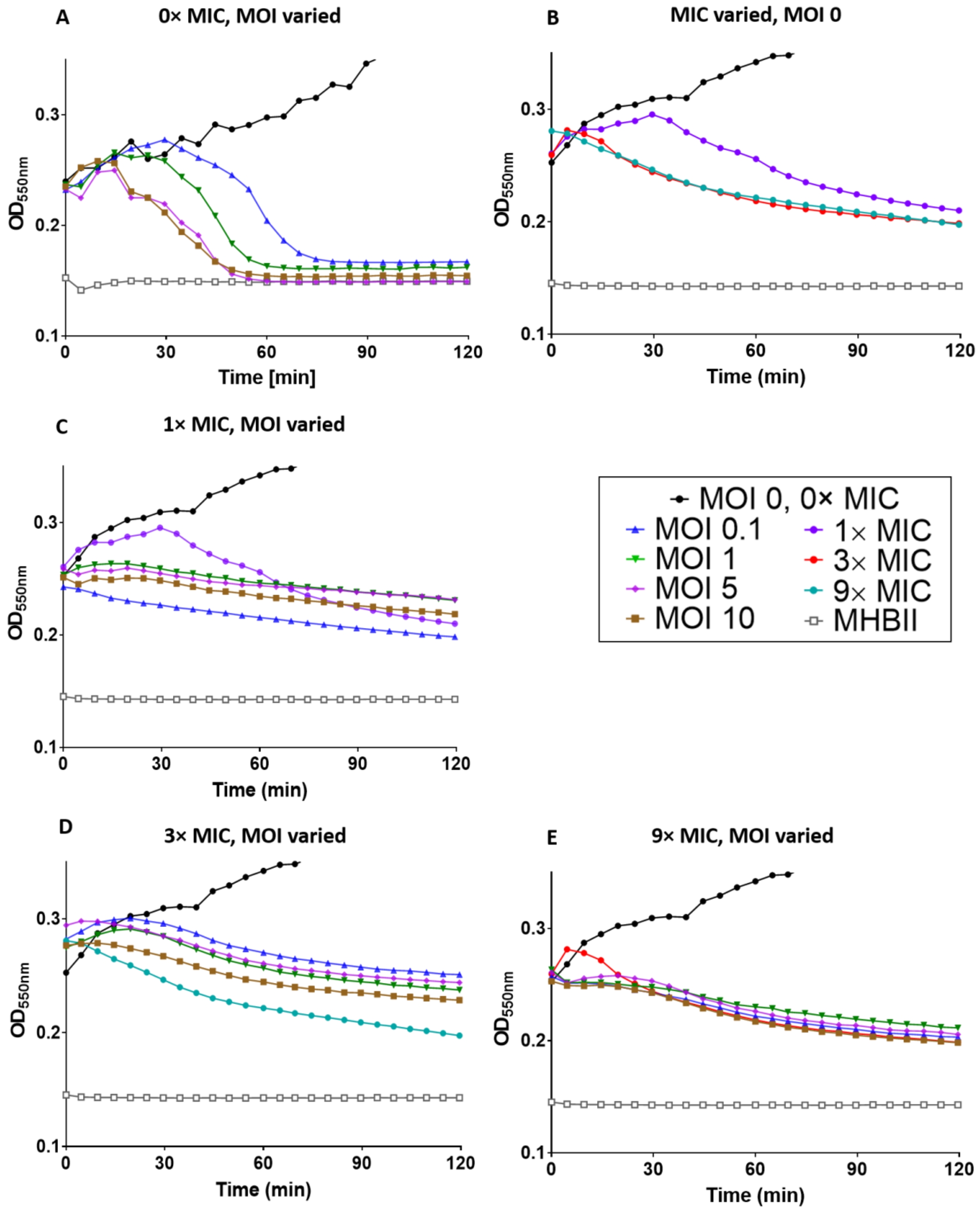


**Figure S2. High antagonism of colistin on ϕKMV infection activities.** See Fig. S1 legend for details. Phage ϕKMV belongs to *phiKMVvirus* and uses type IV pili as a surface receptor.

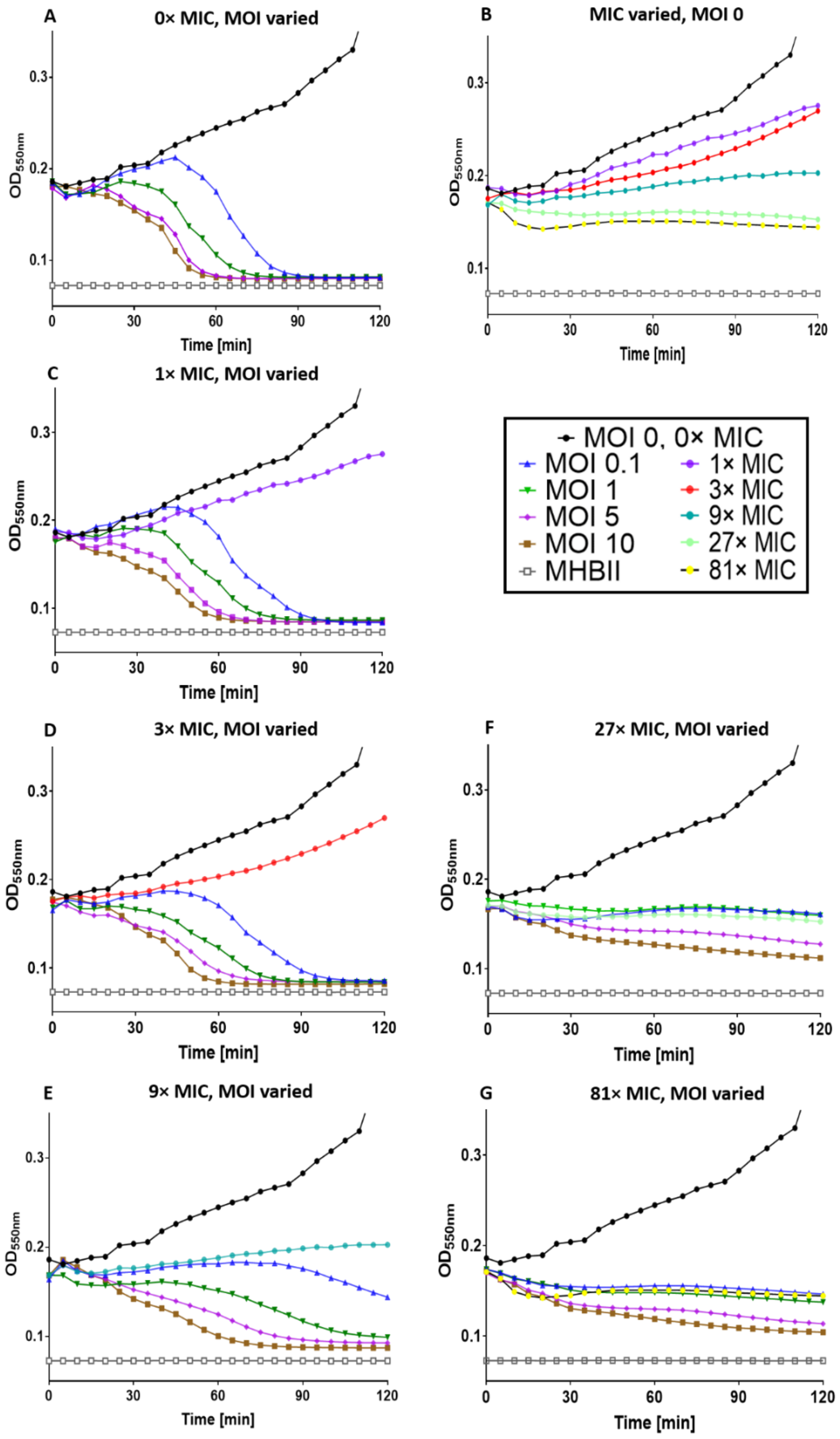


**Figure S3. Low antagonism of ciprofloxacin on LUZ19 infection activities.** See Fig. S1 legend for details.

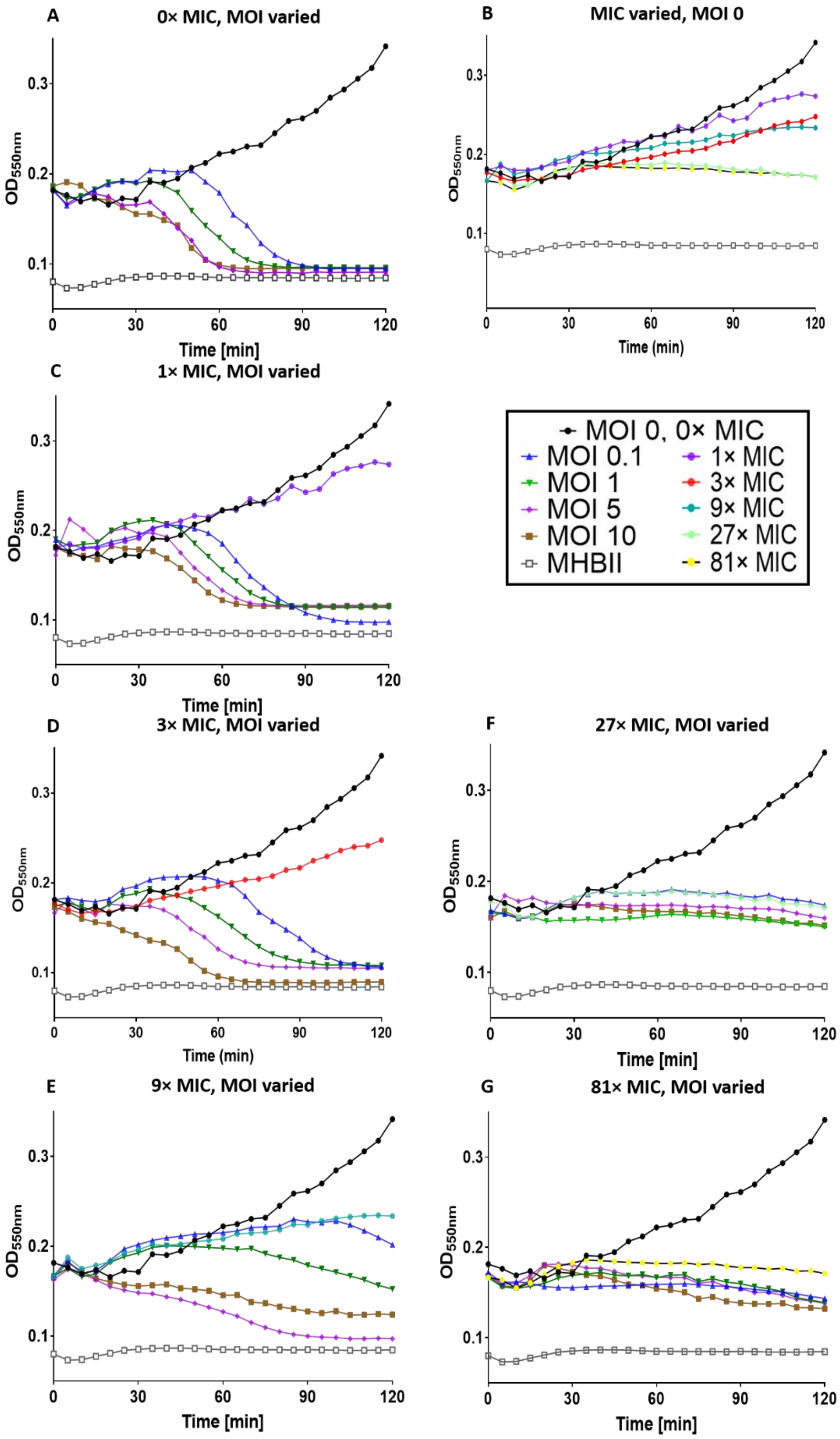


**Figure S4. Low antagonism of ciprofloxacin on ϕKMV infection activities.** See Fig. S1 legend for details.

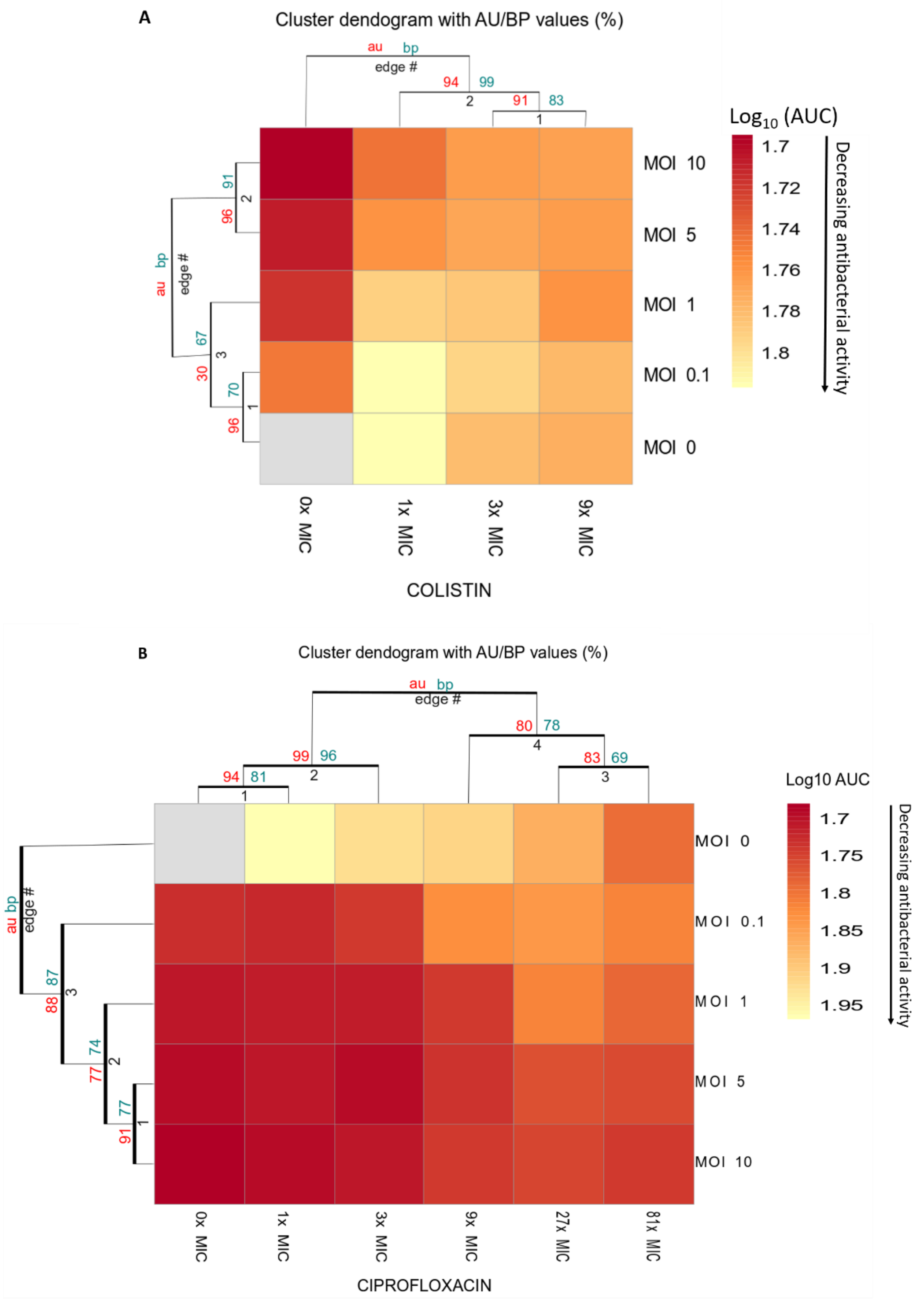


**Figure S5. heat map analyses of PEV2 infection with colistin (A) and ciprofloxacin (B).** Heat maps were created based on $\log_{10}$(AUC). Calculations in each panel are based on 6 replicates from 2 independent experiments. [*] p<0.005, ANOVA; [AU] approximately unbiased *p*-value; [BU] bootstrap probability value, [edge #] *p*-values (%).
